# Using AlphaFold to predict the impact of single mutations on protein stability and function

**DOI:** 10.1101/2021.09.19.460937

**Authors:** Marina A. Pak, Karina A. Markhieva, Mariia S. Novikova, Dmitry S. Petrov, Ilya S. Vorobyev, Ekaterina S. Maksimova, Fyodor A. Kondrashov, Dmitry N. Ivankov

## Abstract

AlphaFold changed the field of structural biology by achieving three-dimensional (3D) structure prediction from protein sequence at experimental quality. The astounding success even led to claims that the protein folding problem is “solved”. However, protein folding problem is more than just structure prediction from sequence. Presently, it is unknown if the AlphaFold-triggered revolution could help to solve other problems related to protein folding. Here we assay the ability of AlphaFold to predict the impact of single mutations on protein stability (ΔΔG) and function. To study the question we extracted metrics from AlphaFold predictions before and after single mutation in a protein and correlated the predicted change with the experimentally known ΔΔG values. Additionally, we correlated the AlphaFold predictions on the impact of a single mutation on structure with a large scale dataset of single mutations in GFP with the experimentally assayed levels of fluorescence. We found a very weak or no correlation between AlphaFold output metrics and change of protein stability or fluorescence. Our results imply that AlphaFold cannot be immediately applied to other problems or applications in protein folding.

## 1 Introduction

AlphaFold is widely claimed to have revolutized protein 3D structure prediction from protein sequence, a 50-years long-standing challenge of protein physics and structural bioinformatics [1]. The fourteenth round of CASP, a blind competition on protein 3D structure prediction [2], demonstrated that AlphaFold, a newcomer to the field, significantly outperforms all other methods. Crucially, AlphaFold models showed an accuracy of their predicted structures that was comparable to structures solved by experimental methods, like X-ray crystallography, NMR, and Cryo-EM [3].

‘It will change everything’, said Andrei Lupas in an interview to Nature [3]. One of the primary changes may be that AlphaFold may also solve other problems related to protein folding. These problems include the prediction of various protein interactions, such as protein-protein, proteinligand and protein-DNA/RNA, and the prediction of the impact of mutations on protein stability. AlphaFold proved to be useful for experimental determination of protein structures with molecular replacement phasing [4, 5] and already facilitated elucidation of SARS-Cov2 protein structures [6, 7]. Furthermore, AlphaFold in collaboration with EMBL-EBI launched a global initiative on constructing the structure models for the whole protein sequence space [8]. The database of freely available structures of all human proteins, and other 20 key organisms yet to be determined, is attributed to “revolutionize the life sciences” [3]. Furthermore, AlphaFold is expected to bring new insights into our understanding of the structural organization of proteins, boost the development of new drugs and vaccines [9]. Researchers in the field are already actively testing AlphaFold performance in various bioinformatics tasks, for instance, in peptide-protein docking [10, 11].

Guided by the expected immediate impact of AlphaFold for the solution of a wide range of problems in structural bioinformatics, we explored the capacity of AlphaFold predictions to serve as a proxy for the impact of mutations on protein stability change (ΔΔG). Although AlphaFold provides a disclaimer that it “has not been validated for predicting the effect of mutations” (https://alphafold.ebi.ac.uk/faq), the expectations of AlphaFold are so high that we judged it prudent to check how well AlphaFold predictions could work for estimation of ΔΔG values. We found that the difference between pLDDT scores, the only local AlphaFold prediction metric reported in the output PDB file, had a very weak correlation with experimentally determined ΔΔG values (Pearson correlation coefficient, PCC = 0.17). The difference in the global AlphaFold metric - the pLDDT averaged for all residues - shows no correlation, both isolated and in combination with the mutated residue’s pLDDT score. Similarly, the same AlphaFold metrics had a very weak correlation with the impact of single mutations on protein function, fluorescence, of GFP. Recent results show that the use of AlphaFold models instead of template structures does not improve ΔΔG prediction by FoldX [12] [13]. Taken together, so far we do not see a use for AlphaFold to help solve the problem of predicting the impact of a mutation on protein stability. The availability of AlphaFold models allows applying more accurate 3D protein structure-based ΔΔG predictors rather than sequence-based ΔΔG predictors; the bottleneck still seems to be the accuracy of current 3D protein structure-based ΔΔG predictors.

## 2 Results

### 2.1 Data set of mutations

We used experimental data on protein stability changes upon single-point variations from ThermoMut Database [14]. After the filtering procedure (see Methods) we randomly chose 976 mutations in 90 proteins for our analysis. For the multiple linear regression analysis, the dataset was split into two sets, a training and a testing set. The split was based on BLAST [15] results, such that the mutations were assigned to the testing set if corresponding proteins had <50% sequence identity to any other protein in the entire dataset (see Methods). All of the other mutations were assigned to the training set.

### 2.2 AlphaFold prediction metrics

Along with coordinates of all heavy atoms for a protein, AlphaFold model contains “its confidence in form of a predicted lDDT-C*α* score (pLDDT) per residue” [1]. LDDT ranges from 0 to 100 and is a superposition-free metric indicating to what extent the protein model reproduces the reference structure [16]. The pLDDT scores averaged across all residues designate the overall confidence for the whole protein chain (<pLDDT>). For each mutation in the dataset, we calculated the difference in pLDDT between the wild type and mutated structures in the mutated position as well as the difference in <pLDDT> between wild type and mutant protein structure models. By checking ΔpLDDT and Δ<pLDDT> values as potential proxies for the change of protein stability we explored the hypothesis that the change of protein stability due to mutation is somehow reflected in the difference of AlphaFold confidence between wild type and mutant structures.

### 2.3 Correlation between ΔΔG and ΔpLDDT values

First, we studied the relationship between the effect of mutation on protein structure stability and the difference in the accuracy of protein structure prediction by AlphaFold for the wild-type and mutant proteins. We did not observe a pronounced correlation between the mutation effect and the difference in confidence metrics (Figure 1). The correlation coefficient is 0.17 ± 0.03 (p-value < 10^-7^) for ΔpLDDT and 0.00 ± 0.03 (p-value = 0.93) for the Δ<pLDDT>.

**Figure 1:**
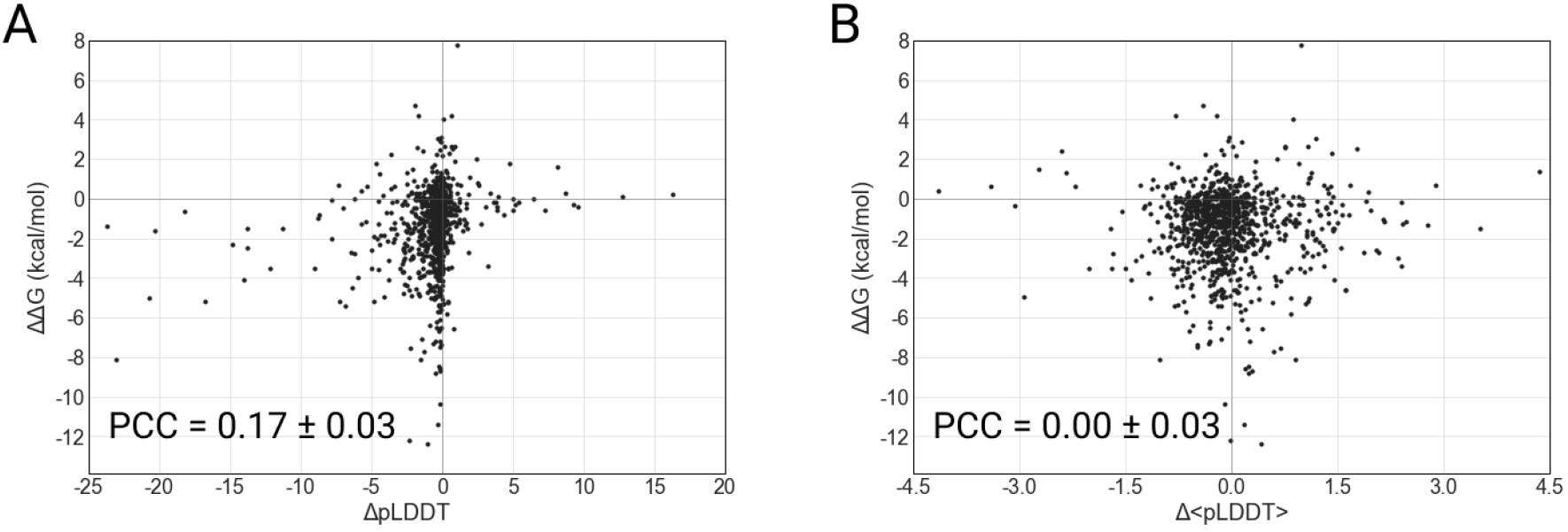
Correlation between the effect of mutation and change of confidence score of structure prediction for the mutated amino acid (A) and for the whole structure (B).

Since the confidence metrics for a given amino acid and whole protein are weakly correlated (PCC = 0.20 ± 0.03, p-value < 10^-9^) we then explored how their combination correlates with the effect of mutation. Multiple linear regression model resulted in the dependence ΔΔG = −1.08 + 0.11·ΔpLDDT + 0.05·Δ<pLDDT>. We did not obtain any pronounced correlation either for training (0.12 ± 0.05, p-value = 0.03) or test sets (0.20 ± 0.04, p-value < 10^-5^).

### 2.4 Relationship of ΔpLDDT and amino acid size change

We explored the outliers with high absolute values of ΔpLDDT in Figure 1A. We noticed the association between ΔpLDDT score and the change of the side chain size upon mutation; to check the association, we conducted the Kruskal-Wallis H-test. The ΔpLDDT scores within different changes in amino acid volume were significantly different (p-value = 0.02). We observed that most of the outlier mutations had a destabilizing effect. We assessed the correlation between the destabilizing effect of a mutation and the decrease in the confidence score. The PCC was greater than for the whole set of mutations, but still not strong (PCC = 0.38 ± 0.08, p-value < 10^-5^). However, the size-changing mutations also populate the region of near-zero pLDDT values, so these mutations also poorly predict ΔΔG from ΔpLDDT (PCC = 0.17 ± 0.07, p-value = 0.01).

### 2.5 Correlation between GFP fluorescence and ΔpLDDT values

Protein stability is intimately coupled with protein functionality. Thus, a reasonable hypothesis holds that the loss of protein functionality due to mutations in most cases results from reduced stability [17]. Therefore, along with testing correlation of AlphaFold metrics with ΔΔG, it is reasonable to test the correlation of AlphaFold metrics with protein function. Furthermore, the change of pLDDT scores may contribute directly to protein functionality without contributing to protein stability. We checked the correlation between ΔpLDDT values and the fluorescent level of 447 randomly chosen single GFP mutants from [18]. The correlation coefficient is 0.14 ± 0.05 (p-value = 0.003) for ΔpLDDT and 0.16 ± 0.05 (p-value < 10^-3^) for the Δ<pLDDT> (Figure 2).

**Figure 2:**
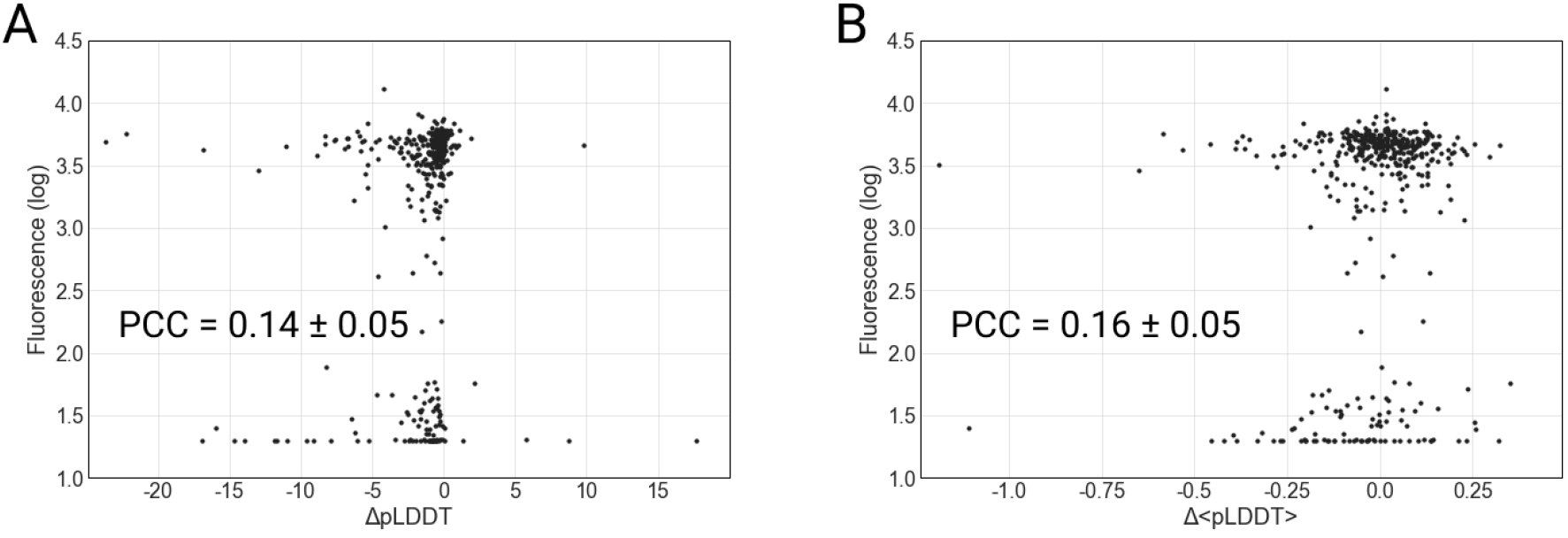
Correlation between the GFP fluorescence and change of confidence score of structure prediction for the mutated amino acid (A) and for the whole structure (B).

## 3 Discussion

Extraordinary success of AlphaFold in predicting protein 3D structure from protein sequence may lead to temptation to apply this tool to other questions in structural bioinformatics. Here we checked the potential of AlphaFold metrics to serve as a predictor for the impact of mutation on protein stability and function. We found a weak correlation of 0.17 ± 0.03 between ΔpLDDT and ΔΔG associated with specific mutations. Although the correlation was statistically significant (p-value < 10^-7^), it is so weak that it cannot be used for accurate ΔΔG predictions (Figure 1) and it is unclear how such predictions can be used in practical applications. Clearly, ΔpLDDT would show a better correlation with ΔΔG if it was measured across bins of averaged ΔΔG. Alternatively, ΔpLDDT could be a separate term in a multiple linear regression model. The averaged metric Δ<pLDDT> shows correlation with ΔΔG, which is statistically indistinguishable from zero. However, a linear combination of the two metrics, ΔpLDDT and Δ<pLDDT>, does not greatly improve the correlation. As for the loss-of-function prediction, the correlation with the impact of mutation on GFP fluorescence showed similar results: PCC was 0.14 ± 0.05 and 0.16 ± 0.05 for ΔpLDDT and Δ<pLDDT>, respectively (Figure 2).

Taken together, our data indicate that AlphaFold predictions cannot be used directly to reliably estimate the impact of mutation on protein stability or function. But why should we have expected such a correlation in the first place? Indeed, AlphaFold was not designed to predict the change of protein stability or function due to mutation. In the words of the authors “AlphaFold is not expected to produce an unfolded protein structure given a sequence containing a destabilising point mutation” (https://alphafold.ebi.ac.uk/faq). However, the only reason for a protein to fold into the distinct native structure is the stability of this structure, so the protein 3D structure and its stability are closely connected. Logically, an algorithm predicting protein 3D structure from sequence should search for the most stable 3D state under the native (or standard) conditions. If a compact structure becomes unstable (for example, due to mutation) then we might expect that the algorithm shifts its predictions toward an unfolded state. Evidence in favor of this point of view is the successful prediction of natively disordered protein regions by AlphaFold and the correlation between the decrease of pLDDT and propensity to be in a disordered region [19]. Thus, it is not unreasonable to expect a decrease in the confidence score of the mutated residue or the whole native structure.

Our results show that AlphaFold repurposing for ΔΔG prediction did not work for the proteins we studied. However, AlphaFold may be used to help predict the impact of a mutation on protein stability or function using AlphaFold 3D models for 3D-structure based ΔΔG predictors. Indeed, it was reported many times that 3D-based predictors perform better than 1D-based [20, 21, 22], so the availability of a pool of high-quality 3D predicted structures could be a plus. However, the performance of the resulting predictions is going to be far from perfect: the 3D-structure based ΔΔG predictors are far from being perfect even using 3D structures from PDB [23], they show correlation of 0.59 or less in independent tests [24]. Thus, using AlphaFold models instead of PDB structures does not make ΔΔG predictions more accurate [13], so availability of AlphaFold models is expected to show an approximately 0.59 correlation with predictions of ΔΔG, which may be too low for many applications.

Will the newly AlphaFold-generated 3D structures for proteins with unknown 3D structures be useful for the training of the new ΔΔG predictors? It seems unlikely, because the vast majority of the current experimental ΔΔG values relate to proteins with known 3D structures. Thus, the newly generated structures will be useful to the same extent as the entire PDB has been across the last few decades.

The deep learning approach demonstrated by AlphaFold may be an inspiring example to develop a deep learning ΔΔG predictor. However, we see the dramatic difference between the situations with 3D structure prediction and ΔΔG prediction that may impede this development. The difference is in the amount of available data. For protein structure prediction AlphaFold used PDB with >180,000 files, and each file contained a wealth of information. In contrast to PDB, the number of experimentally measured ΔΔG values are of the order of 10,000 and these are just numbers without accompanying extra data. To make a rough comparison of information in bits, PDB structures occupy 100 Gb, while all the known experimentally ΔΔG values occupy 10 kb. Neural networks are very sensitive to the amount of information in the training set so the ability of deep learning to tackle the ΔΔG prediction task at present looks hindered mostly by the lack of experimental data.

Overall, we explored the capacity of direct prediction of ΔΔG by all of the reported AlphaFold metrics: (i) the difference in the pLDDT score before and after mutation in the mutated position, (ii) the difference in the averaged pLDDT score across all positions before and after mutation. We found that the correlation was weak or absent, and, therefore, AlphaFold predictions are unlikely to be useful for ΔΔG predictions. Taken together with our recent result that AlphaFold models are not better for ΔΔG predictions by FoldX than best templates [13], we see no straightforward way to use AlphaFold advances for solving the task of prediction of ΔΔG upon mutation. The task of ΔΔG prediction should be solved separately and it will face the problem of limited amount of data for training neural networks.

## 4 Methods

### 4.1 Dataset of experimental mutations

The data on experimentally measured effects of mutations on protein stability were taken from ThermomutDB [14] (version 1.3). From 13,337 mutations in the database we extracted single-point mutations with data on ΔΔG.

The filtered dataset resulted in 5712 mutations in 286 proteins. We have done the analysis for randomly chosen 976 mutations in 90 proteins.

### 4.2 Dataset of GFP mutants fluorescence

We took data on fluorescence levels of GFP mutants from [18]. From the original dataset we randomly extracted 447 single mutants for our analysis.

### 4.3 Protein structure modeling with AlphaFold

The wild type protein structures were retrieved from the AlphaFold Protein Structure Database (AlphaFold DB) [8] by their Uniprot accession code. The structures of original proteins that were absent in the AlphaFold DB as well as structures of mutant proteins were modeled by the standalone version of AlphaFold [1] using the fasta file with Uniprot sequence of a protein as the only input in the ‘–fasta_paths’ flag.

### 4.4 Prediction metrics

The per-residue local distance difference test (pLDDT) confidence scores for the protein structure models downloaded from the AlphaFold DB were retrieved from the B-factor field of the coordinate section of the pdb file. The pLDDT confidence scores for the protein structure models that we predicted by standalone AlphaFold were extracted from the pickle file for the best ranked model from “plddt” array.

### 4.5 Sequence identity of proteins within the dataset of mutations

To identify the sequence identities of the proteins in the dataset of mutations we performed protein BLAST [15] search of protein sequences against themselves.

We divided the dataset into training and test sets for linear regression model based on the arbitrary sequence identity threshold of 50%. Mutations in proteins above the threshold comprised the training set, and the rest of mutations were used as the test set. The training and test sets resulted in 364 mutations in 60 proteins and 612 mutations in 30 proteins.

### 4.6 Linear regression analysis

Multiple linear regression fit with two parameters was performed using the linear_model module of Sklearn library with default parameters.

## 5 Acknowledgments

The authors thank Zimin Foundation and Petrovax for support of the presented study at the School of Molecular and Theoretical Biology 2021. The authors acknowledge the use of Zhores supercomputer [25] for obtaining the results presented in this paper.

## Abbreviations

3D: three-dimensional
CASP: critical assessment of protein structure prediction
PDB: protein data bank
NMR: nuclear magnetic resonance
DNA: deoxyribonucleic acid
RNA: ribonucleic acid
PCC: Pearson correlation coefficient
LDDT: local distance difference test
pLDDT: per-residue local distance difference test
GFP: green fluorescence protein
BLAST: basic local alignment search tool

